# Redundant *PLETHORA* activity promotes development of early embryonic cell lineages in Arabidopsis

**DOI:** 10.1101/2022.03.02.482431

**Authors:** Merijn Kerstens, Carla Galinha, Hugo Hofhuis, Michael Nodine, Ben Scheres, Viola Willemsen

## Abstract

The BABY BOOM/PLETHORA4 (BBM/PLT4) transcription factor has received much attention due to its ability to induce somatic and zygotic embryogenesis, two processes of pivotal importance in plant breeding. Loss of additional *AINTEGUMENTA-LIKE/PLETHORA* (*AIL/PLT*) genes, encoding members of the APETALA2 transcription factor family, causes embryo arrest and abortion, but whether BBM/PLT4 provides specific information for embryo development has remained unknown. Here, we reveal that *AIL/PLT* members are expressed in partially overlapping domains from their first appearance in the apical cell daughter of the zygote. Redundant early embryonic activity of BABY BOOM/PLT4 and PLT2 triggers development of the apical cell lineage and is required to initiate embryonic primordia. Furthermore, promoter swap experiments show that *PLT1* and *PLT3* expression in the expression domains of *PLT2* and *BABY BOOM/PLT4* is sufficient to rescue *plt2 bbm* double mutants. Our data indicate that generic AIL/PLT factors, involved in maintenance of stem cells, promotion of cell division and suppression of cell differentiation, provide the necessary information to initiate embryogenesis in Arabidopsis.

## Introduction

After fertilization in flowering plants, the zygote expands and cell division as well as pattern formation are initiated to ultimately generate the mature embryo. The embryo is surrounded by the endosperm, resulting from a second fertilization of the central cell, and maternally derived seed tissues. It is well established that ectopic expression of the AINTEGUMENTA-LIKE/PLETHORA (AIL/PLT) clade transcription factor BABY BOOM/PLETHORA4 (BBM/PLT4) can initiate somatic embryos (Boutilier et al., 2002; Horstman et al., 2017; Tsuwamoto et al., 2010), an agronomically relevant feature as somatic embryogenesis can support regeneration processes in many breeding applications. In addition, induction of *BBM*-like genes in the egg cell can trigger asexual embryogenesis (Conner et al., 2015; Conner et al., 2017; El Ouakfaoui et al., 2010). In rice, ectopic egg cell expression substitutes for the absence of paternally derived BBM1 which, together with mutations that substitute meiosis for mitosis (Khanday et al., 2019), can be used for clonal propagation with immense value in breeding and seed production.

PLETHORA (PLT) transcription factors of the APETALA2/ETHYLENE-RESPONSIVE ELEMENT BINDING PROTEIN family are key post-embryonic developmental patterners (Horstman et al., 2014; Scheres and Krizek, 2018). In roots, PLTs indirectly read out an auxin gradient to establish stable tissue zonation by controlling cell division and differentiation (Mahonen et al., 2014; Roszak et al., 2021; Santuari et al., 2016) and they are required for stem cell maintenance (Aida et al., 2004; Blilou et al., 2005). Although PLTs are involved in a wide array of developmental processes including development of the primary roots (Aida et al., 2004; Galinha et al., 2007), lateral roots (Du and Scheres, 2017; Hofhuis et al., 2013; Kerstens et al., 2021), phyllotaxis (Pinon et al., 2013; Prasad et al., 2011), and pluripotency acquisition in callus (Zhai and Xu, 2021), their role during early embryogenesis has not been resolved. However, homozygous combinations of *plt2* and *bbm* in *PLT1, PLT2* seedlings were not recovered when we attempted to generate higher-order mutants of, *AIL/PLT* clade members (Galinha et al., 2007), suggesting that specific PLTs are required for early embryogenesis. In support of this notion, ectopic overexpression of PLTs promoted the production of embryos from vegetative tissues (i.e. somatic embryogenesis). We therefore investigated which *PLT* genes are functional at the initiation of embryogenesis to define primordial cells that are precursors to all embryo cell types.

## Results

### *PLTs* are expressed in overlapping domains throughout embryogenesis

To determine *PLT* expression patterns during early embryogenesis, we examined functional PLTx:YFP translational fusions under the control of their native promoters (pPLTx::PLTx:YFP; (Aida et al., 2004; Galinha et al., 2007) from the one-cell stage onward. We detected *PLT2:YFP, BBM:YFP* and *PLT7:YFP* expression in both the apical and basal cells of one-cell stage embryos; *PLT1:YFP* was only expressed in the basal cell (Fig. 1A). *PLT3:YFP* and *PLT5:YFP* were not detectable at this stage. At the two/four-cell stage, *PLT2, BBM* and *PLT7* remained expressed in the apical cells and basal cell, whereas *PLT5* expression was weakly visible in the apical cells and basal cells (Fig. 1B). *PLT3* could not be detected at this point in time. From the globular stage onward, all *PLTs* were expressed in the embryo proper (Figs. 1C; S1). In contrast to *PLT3* and *BBM*, which were expressed in all cells of the globular-stage embryo proper, the expression of *PLT1* and *PLT2* was restricted to the lower tier of the proembryo (Fig. 1C). In a similar fashion, PLT5 was specifically restricted to the upper tier from the globular stage to the triangular stage (Figs. 1C; S1). *PLT7* expression was localised in the outer layer (the protoderm) of the proembryo (Fig. 1C). At the heart stage, expression of *PLT1, PLT2, PLT3* and *BBM* extended to the quiescent centre (QC) progenitor lens-shaped cell (Fig. S1). In later stages, these four PLTs were detected in the QC, columella and in the vascular tissue (Fig. S1). Taken together, *PLT* expression patterns form distinct overlapping domains throughout embryogenesis (Fig. 1D).

**Figure 1.**
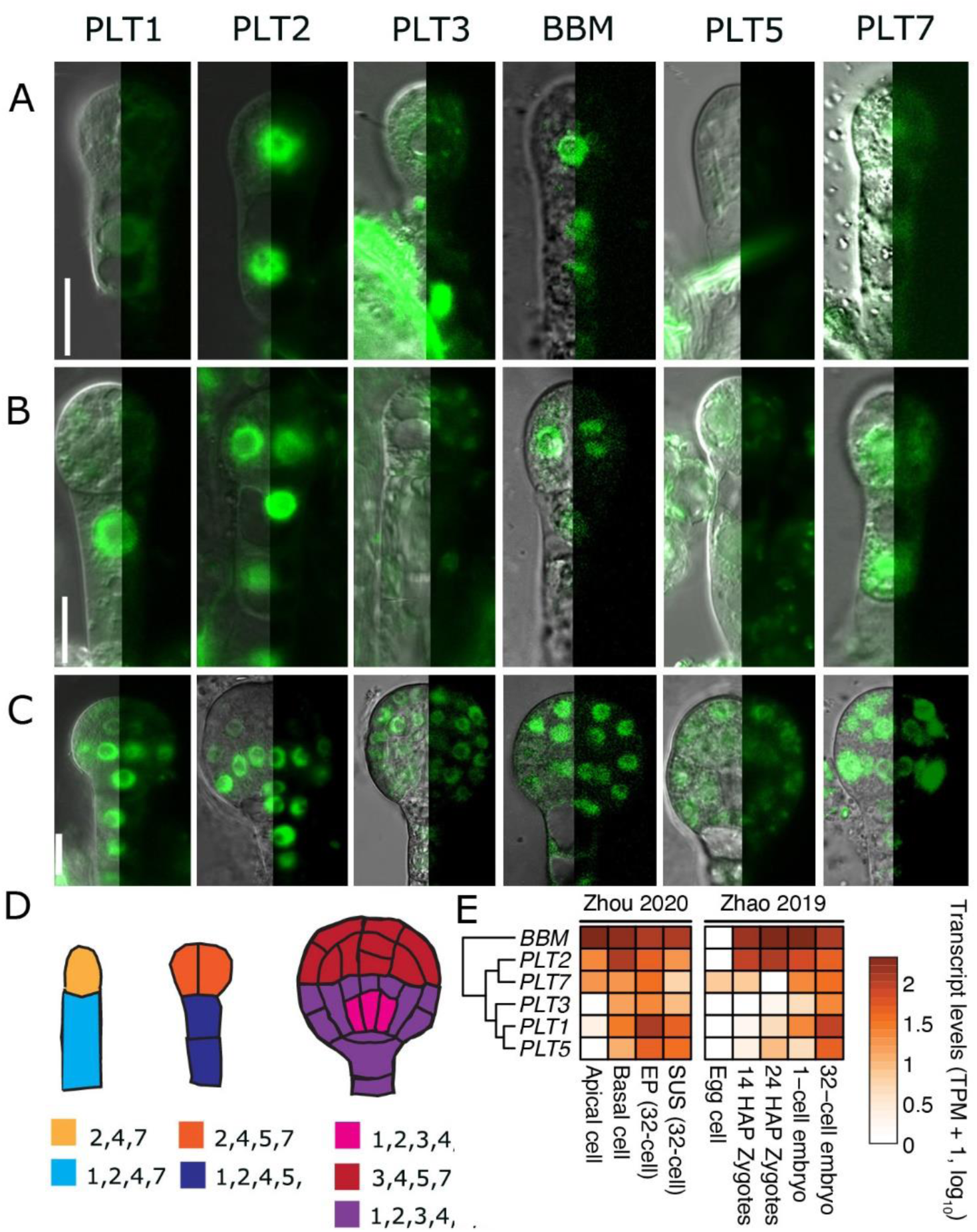
*PLT* genes are expressed in partially overlapping expression domains in early embryos. pPLTx::PLTx:YFP signal in one-cell stage embryos (A), two-to-four-cell stage embryos (B), and globular embryos (C). DIC and confocal signal are shown per image. Scale bars correspond to 10 μm (D) Schematic representations of *PLT* expression domains per stage is marked with different colours; numbers denote PLT identity (e.g. 1 = PLT1, 4 = BBM/PLT4 (E) PLTs are hierarchically clustered according to transcripts per million (TPM) values obtained from Zhou et al. (2020). EP = embryo proper, SUS = suspensor, HAP = hours after pollination.

In our expression analysis, we deliberately chose to avoid transcriptional fusions to retain potential regulatory elements in intronic sequences. To gain insight in the native transcriptional dynamics of *PLTs* during early embryogenesis, we mined three RNA-seq datasets of early embryos (Hofmann et al., 2019; Zhao et al., 2019; Zhou et al., 2020). In agreement with our observations from the translational fusions, *PLT1, PLT2, BBM* and *PLT7* were expressed moderately to strongly in contrast to the weakly expressed *PLT3* and *PLT5* at the one-cell stage (Fig. 1A,E). Importantly, *PLT2, BBM* and *PLT7* (and to a minor degree *PLT1*) were the only *PLTs* expressed in the apical cell at this stage, reinforcing their role in early embryogenesis (Fig. 1E). From the 32-cell stage onward, all *PLTs* were expressed in embryos (Figs. 1E; S2), which was consistent with our translational reporters. The localisation patterns observed at the globular stage also largely overlapped those obtained by single-nucleus mRNA-sequencing of globular embryos ((Kao et al., 2021);Fig. S3). We thus demonstrate that our translational YFP fusions are in strong congruence with published transcriptome data and that multiple *PLTs* are expressed during early embryogenesis.

### PLT2 and BBM are essential for early embryonic cell progression

Conservation of PLT sequence, copy number and genomic context within and between angiosperm lineages (Kerstens et al., 2020), as well as their overlapping expression patterns, suggests redundant roles during embryogenesis. Since *plt2 bbm* double mutants could not be recovered previously (Galinha et al., 2007) and the expression domains of *PLT2* and *BBM* strongly overlapped in one-cell stage embryos, we generated *plt2-2/+ bbm-1* (hereafter *plt2/+ bbm*) mutants to inspect early embryo development of the selfed progeny (Fig. 2A). In the analysis of the offspring, we found that approximately ∼25% of the seeds arrested development (Fig. 2B; Table 1). In wild type plants, the embryo could be easily recognized within the ovule. In *plt2/+ bbm* plants, however, embryos were inconspicuous in ∼25% of the ovules (Table 1). A closer examination of these ovules revealed that the embryos arrested development at a very early stage. Arrested embryos were most easily recognized when the non-arrested embryos in the same silique were at globular stage (Fig. 2Ci). Embryos arrested before or at the one-cell stage; the zygote appeared to fail to elongate in 69% (20/29) of these apparently empty ovules, and after the first cell division development stopped in 24% (7/29) while in 7% (2/29) the basal cell was still able to divide (Fig. 2Cii-iv). The frequency of arrested embryos is as expected for a fully recessive *bbm* allele, corroborating that the molecular lesion conferred a loss-of-function phenotype. Furthermore, this frequency indicates that the effect of the mutations occurs post-fertilization rather than being a maternal effect.

**Table 1.** Aborted and non-aborted embryos in higher-order *plt* mutants.

| Genotype | Early-aborted embryos (#) | Non-aborted embryos (#) | Early-aborted embryos (%) |
| --- | --- | --- | --- |
| *plt2/+ bbm* | 526 | 1958 | 26.9 |
| *plt2 plt7* | 0 | 350 | 0.0 |
| *bbm plt7* | 0 | 234 | 0.0 |
| *plt3 bbm plt5* | 0 | 1200 | 0.0 |
| *plt1 plt2 plt3 plt5* | 0 | 300 | 0.0 |
| *plt1 plt2 plt3 plt7* | 0 | 570 | 0.0 |
| *plt3 bbm plt5 plt7-cr* | 1 | 148 | 0.7 |

**Figure 2.**
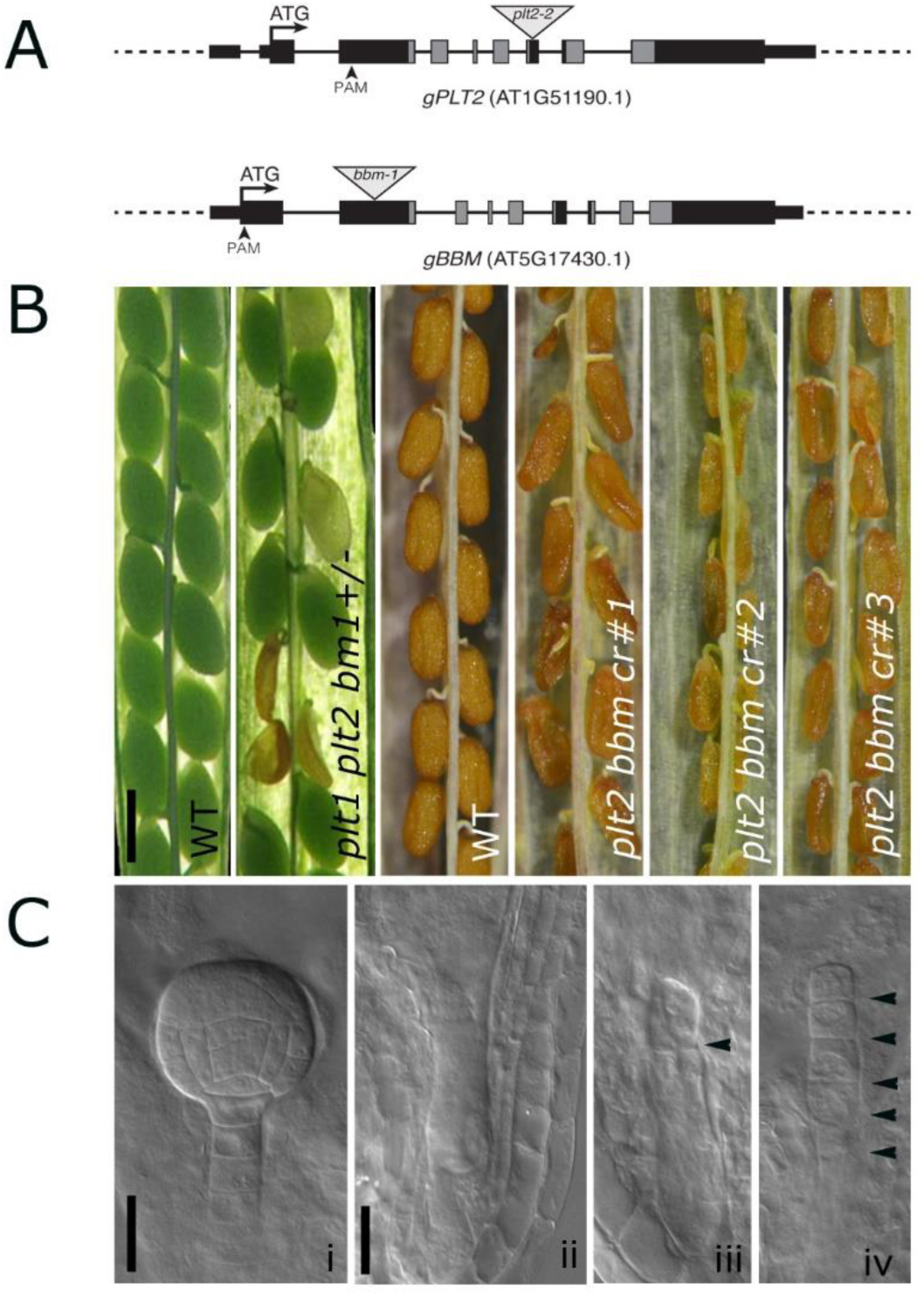
PLT2 and BBM are required for early embryogenesis. (A) Gene models of *PLT2* (AT1G51190.1) and *BBM* (AT5G17430.1). Black rectangles and lines indicate exons and introns, respectively. UTRs are denoted by thin rectangles. AP2 domains are marked in grey. Triangles denote T-DNA insertion sites. Arrowheads indicate CRISPR PAM sites. (B) Aborted seeds and viable seeds in siliques of WT and *plt1 plt2 bbm/+* self-fertilized plants and in T_1_ lines with CRISPR mutations (*plt2 bbm-cr*) and without (WT), respectively. (C) Phenotype of embryos from segregating *plt2 bbm/+* plants at comparable stage of development, (i) normal globular stage embryo, (ii-iv) arrested embryos. Arrow heads indicate division planes. Scale bar in panel B corresponds to 0.5 mm and in panel C to 50 μm.

We next extended our analysis and investigated whether other family PLT members, in particular the early-expressed *PLT7*, are redundantly required during one-cell stage development. Although *PLT7* is expressed at the one-cell stage in the apical and basal cells, we did not identify embryonic abortion in *plt2 plt7* and *bbm plt7* double mutants nor in *plt1 plt2 plt3 plt7* quadruple mutants (Table 1). Since single and higher-order mutants of *plt3, plt5* and *plt7* have been extensively studied for their role in root and shoot architecture; (Du and Scheres, 2017; Hofhuis et al., 2013; Kerstens et al., 2021; Pinon et al., 2013; Prasad et al., 2011), this excludes the possibility that they are mandatory in early embryo development. Moreover, embryogenesis proceeded normally in *plt3 plt5 plt7-cr* mutants (Kerstens et al., 2021) in which we also knocked out *BBM* with CRISPR/Cas9 (*plt3 bbm plt5 plt7-cr*, Table 1). The same holds true for other PLT mutation combinations: early embryos from self-fertilized *plt3 bbm-2* and *plt1 plt3 bbm-2* plants develop without defects (Galinha et al., 2007), as do early embryos of genotypes *plt1 plt2 plt3 plt5, plt3 bbm-1 plt5*, and *plt1 plt2/+ plt3 bbm-2* (Table 1; (Galinha et al., 2007)). Taken together, these results suggest that *BBM* and *PLT2* have a specific role in early development that is not shared by other *PLT* genes.

To exclude that the described phenotype is specific to *plt2* and *bbm* alleles generated by the T-DNA insertions in their coding sequences or the mixed ecotype background constituting the seedlings (Wassilewskija x Col-0), we targeted both genes with a single-guide RNA (sgRNA) to generate frame-shift mutations by CRISPR/Cas9 mutagenesis in Col-0 (Fig. 2A). To circumvent the embryo-lethality phenotype, we directly analysed T_1_ plants with mutations in both *PLT2* and *BBM*, which could germinate from non-double-homozygous seeds after transformation. In three independent T_1_ plants with inflorescences harbouring a different set of homozygous, frame-shift causing insertions in both *PLT2* and *BBM* (*plt2 bbm-cr*), all seeds were aborted (i.e. concave in shape), which did not occur in T_1_ plants without CRISPR mutations in the inflorescences (Fig. 2B). This finding indicates that knocking out *PLT2* and *BBM* results in an arrest at early embryogenesis independent of the genetic background.

### PLT promoter swaps can rescue the *plt2 bbm* phenotype

Since PLT2 and BBM act in a redundant fashion, we asked whether PLT protein identity or spatiotemporal control of *PLT* expression was more important for progression of embryogenesis. To explore whether *PLT2* and *BBM* are interchangeable, pPLT2::BBM:YFP and pBBM::PLT2:YFP promoter swap constructs were introduced in the *plt2/+ bbm* background. Siliques of *plt2/+ bbm* homozygous for the swap constructs were analysed by counting the number of normal and aborted embryos. Both swaps could almost fully rescue the *plt2/+ bbm* embryo phenotype (Table 2), indicating that the *PLT2* and *BBM* expression domains show functional overlap. To investigate whether *plt2/+ bbm* complementation with other *PLTs* driven by *pPLT2* or *pBBM* could be established, we reintroduced *PLT1* and *PLT3*, which are hardly - if at all - expressed in the apical cell of the one-cell stage embryo (Fig. 1A,E). *plt2/+ bbm* mutants were transformed with pPLT2::PLT1:YFP, pPLT2::PLT3:YFP and pBBM::PLT1:YFP to determine the proportion of aborted ovules. All promoter swaps could rescue the one-cell stage defect (Table 2). Thus, despite the tight regulation and activation of *PLT2* and *BBM* during early embryogenesis, reactivation of other PLT family members at this early stage can complement the morphological defects of *plt2 bbm* mutant embryos. Therefore, we conclude that the role of PLT2 and BBM in initiation of embryo development from the one-cell stage onward can be exerted by at least two other PLT family members normally not expressed at this stage.

**Table 2.** Aborted and non-aborted embryos in selfed *plt2/+ bbm* complemented with *PLT:YFP* fusions.

| Construct in selfed *plt2/+ bbm* | Early-aborted embryos (#) | Non-aborted embryos (#) | Early-aborted embryos (%) |
| --- | --- | --- | --- |
| pPLT2::gPLT1:YFP | 10 | 794 | 1.3 |
| pPLT2::gBBM:YFP | 65 | 3666 | 1.8 |
| pPLT2::PLT3:YFP | 0 | 595 | 0.0 |
| pBBM::gPLT2:YFP | 15 | 1009 | 1.5 |
| pBBM::gPLT1:YFP | 16 | 522 | 3.1 |

## Discussion

Here, we demonstrate that *PLT2* and *BBM* act redundantly to promote the transition of the zygote to a developing embryo. We show that these two genes are expressed in the zygote shortly after fertilization and, together with *PLT7*, are expressed in the apical cell after the first asymmetric cell division (Fig. 1). Absence of both *PLT2* and *BBM* leads to embryonic arrest at the zygote/one-cell stage, a phenotype that could not be recapitulated with any other combination of *plt* single and higher-order mutants (Fig. 2; Table 1). Finally, promoter swap experiments demonstrated that spatiotemporal control rather than PLT sequence identity is causal for the observed phenotype (Table 2).

It has been thoroughly established that PLTs orchestrate a wide array of developmental processes (Horstman et al., 2014; Scheres and Krizek, 2018). Given that ectopic overexpression of all six *PLTs* causes production of somatic embryos in Arabidopsis (Boutilier et al., 2002; Horstman et al., 2017; Tsuwamoto et al., 2010), it is not surprising that PLTs are also responsible for proper progression of embryogenesis. Importantly, research in other systems corroborates the critical nature of AIL/PLTs in this process. In rice, it was shown that ectopic overexpression of *OsBBM1* resulted in production of somatic embryos, and that seed viability of *osbbm1 osbbm2 osbbm3* triple mutants was severely compromised (Khanday et al., 2019). Understanding which AIL/PLTs guide early embryogenesis is vital in understanding how different transcriptional networks are employed to facilitate morphogenesis.

From an evolutionary perspective, it is intriguing to observe redundancy of PLT2 and BBM, but not PLT1. PLT1 and PLT2 act redundantly to maintain the root apical meristem after germination in Arabidopsis (Aida et al., 2004; Galinha et al., 2007), but this redundancy is clearly not reflected in embryogenesis. *PLT1* and *PLT2* belong to the same gene clade within the euANT family, which exhibits little copy number variation as well as strongly conserved synteny across eudicots (Kerstens et al., 2020). How such similar paralogs developed differential spatiotemporal regulation in early embryos remains enigmatic, although differential expression was also observed in lateral root primordia (Du and Scheres, 2017). Identifying the regulatory regions responsible can shed light on the factors responsible for early embryonic gene expression of the PLT clade members.

As we have shown through promoter swap assays, it is not PLT protein sequence but rather the expression patterns that reflect analogous gene function. In support of this notion, it was shown that PLTs regulate a common set of target genes and that they all recognize an ANT-like consensus motif (Horstman et al., 2015; O’Malley et al., 2016; Santuari et al., 2016; Yano et al., 2009). Any PLT could at least partially rescue the *plt1 plt2* phenotype when driven by *pPLT2* (Galinha et al., 2007; Santuari et al., 2016), and a full complementation of the *plt3 plt5 plt7* lateral root phenotype was established when any *PLT* was expressed from the primordium-specific promoter *pPLT7-1*.*5kb* (Du and Scheres, 2017). Thus predicting PLT function in other plant species based on orthologous relationships demands caution.

In root meristems, AIL/PLT target gene analysis suggest that these factors promote stem cell maintenance, cell division and growth (Santuari et al., 2016), and repress differentiation (Roszak et al., 2021). The redundant and interchangeable roles of *AIL*/*PLT* genes in early embryogenesis lend support to the scenario that the initiation and maintenance of cell division and growth in embryos is mechanistically linked to meristematic programs. It is therefore plausible that the evolution of a meristematic *AIL/PLT* growth control module predated and was adapted for embryonic growth in the context of seed development.

## Materials and Methods

### Plant material

Arabidopsis thaliana ecotypes Columbia-0 (Col-0) and Wassilewskija (WS) were used. *plt1-4* and *plt2-2* alleles were described in Aida et al., (2004) and *plt3-1* in Galinha et al., (2007). *bbm-1* (SALK_ 097021) was provided by the SALK Institute (Alonso et al., 2003). *plt5-2* and *plt7-1* were described in Prasad et al. (2011). *plt3 plt5 plt7-cr* triple mutant was described in Kerstens et al. (2021). Growth conditions of plants are described in (Willemsen et al., 1998). Translational pPLTx::PLTx:YFP were described in Aida et al. (2004), Galinha et al. (2007), and Prasad et al. (2011).

### Cloning

The promoter swaps were generated in the gateway vector pGreenII 227 by using the promoters and genomic sequences of the translational fusions as described in Aida et al. (2004) and Galinha et al. (2007). *plt2* and *bbm* CRISPR constructs were generated as described in Kerstens et al. (2021), except that we used a *Zea mays* codon-optimised intronised Cas9 protein (zCas9i; Grützner et al., 2021), using one sgRNA per gene (5’-CTTAGGAGTGAGCAAATCGG-3’ and 5’-AACTCGATGAATAACTGGTT-3’, respectively). The zCas9i module pRPS5A::ΩTMV:zCas9i:nosT was assembled by combining pICH41233-pRPS5A (Kerstens et al., 2021), pICH41402 (Addgene #50285; ΩTMV 5’UTR), pAGM47523 (Addgene #153221; zCas9i) and pICH41421 (Addgene #50339; nosT) in pICH47742 (Addgene #48001) using golden gate cloning (Engler et al., 2014). All constructs were electroporated into *Agrobacterium tumefaciens* (C58C1.pMP90) and transformed into Col-0 wild-type plants as described in (Clough and Bent, 1998).

### Mutant genotyping

The T-DNA insertions in *plt1-4, plt2-2, plt3-1, bbm-1, plt5*-*2* and *plt7-1* lines were verified by genotyping as described in Aida et al. (2004), Galinha et al. (2007) and Prasad et al. (2011). Mutant combinations were generated by crossing and lines of different allelic combinations were selected from F2 populations by genotyping. *plt2* and *bbm* CRISPR target regions were amplified with PLT2 CRISPR F (5’-GTTTGCAGCCATACTTGGAG-3’), PLT2 CRISPR R (5’-CTTTCACAGTGGCGACTTCT-3’), BBM CRISPR F (5’-GCCTCGGAAGAAATGAACAT-3’) and BBM CRISPR R (5’-CACAAACCTCGGGAGTGACT-3’) and Sanger-sequenced with either primer. The *plt2 bbm* CRISPR mutants were genotyped in the T_1_ generation; the *plt3 bbm plt5 plt7-cr* mutants were genotyped in the T_3_ generation after crossing out the transgene.

### Microscopy and FP detection

For light microscopy, ovules were collected from developing siliques, cleared and mounted according to Willemsen et al. (1998). For confocal microscopy, dissected embryos were mounted in 5% glucose. YFP was detected with an excitation wavelength of 488 nm.

## Supplemental figures

**Figure S1.**
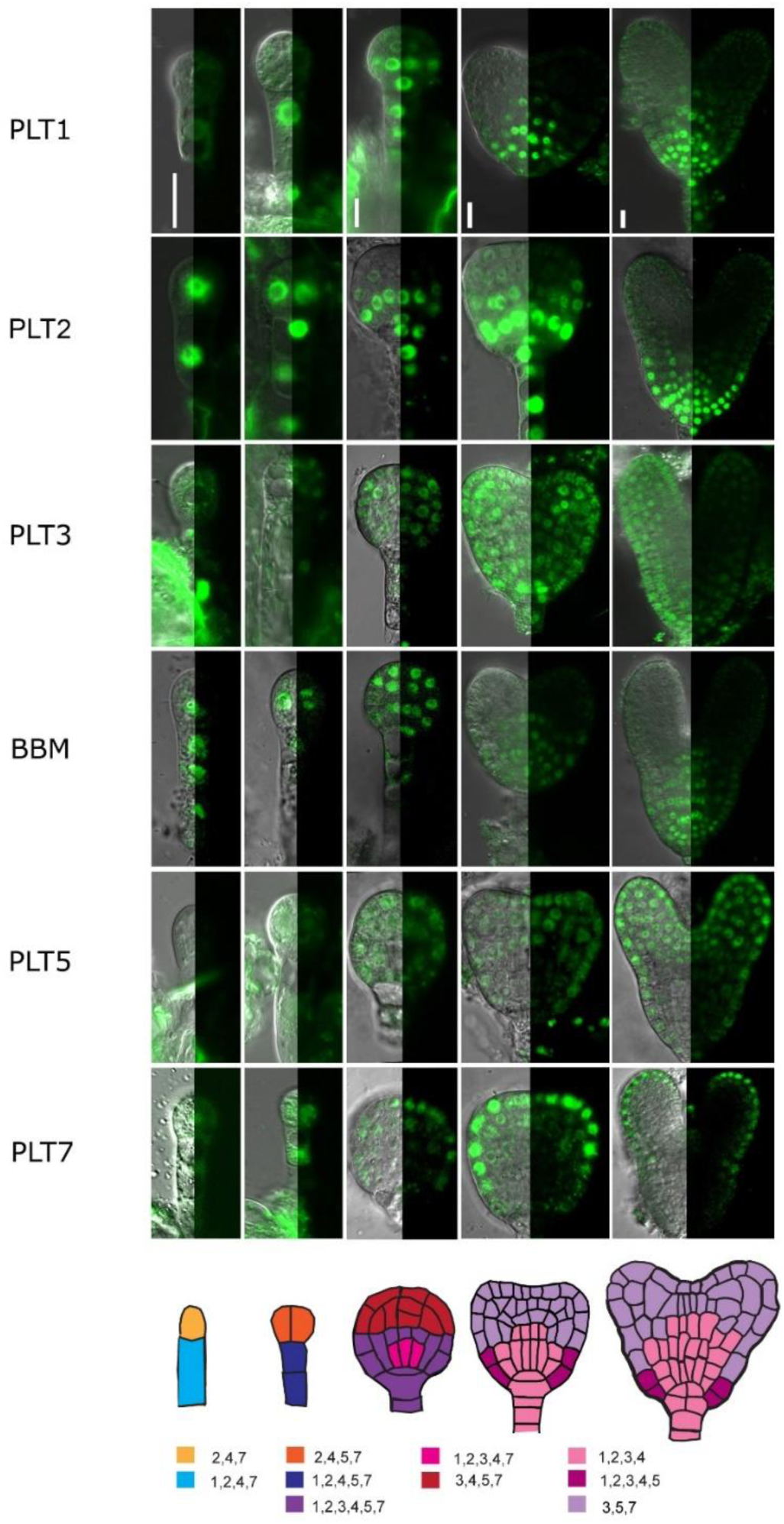
*PLT* genes are expressed in partially overlapping expression domains in late embryos. pPLTx::PLTx:YFP signal in triangular-stage embryos (A), and heart-stage embryos (B). DIC and confocal signal are shown per image. Schematic representations of *PLT* expression domains per stage is marked with different colours; numbers denote PLT identity (e.g. 1 = PLT1, 4 = BBM/PLT4). Scale bars correspond to 10 μm.

**Figure S2.**
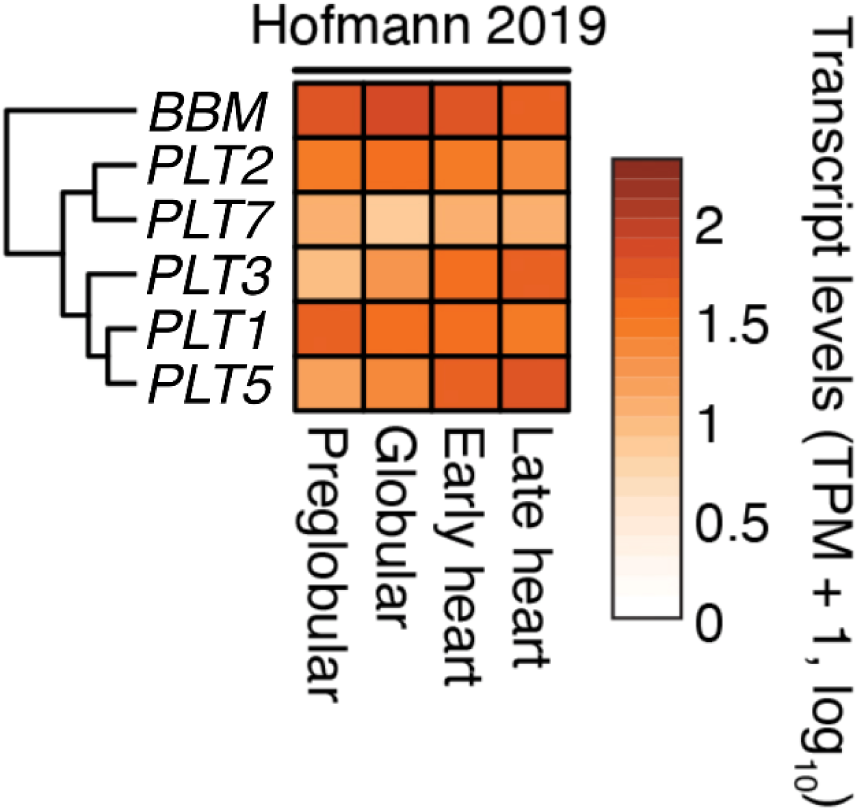
*PLT* expression in preglobular to heart stage embryos (Hofmann et al., 2019). PLTs are clustered according to expression data of Zhou et al. (2020) as displayed in Fig. 2.

**Figure S3.**
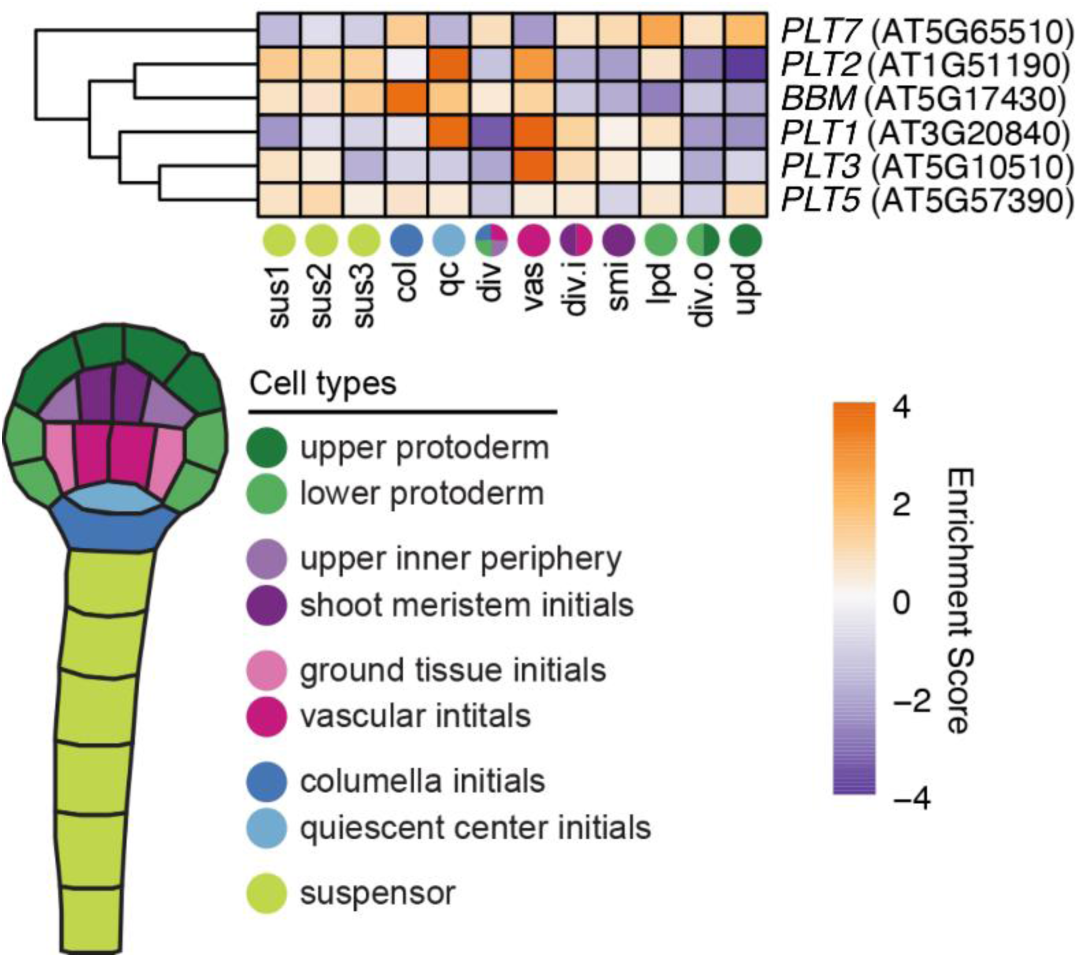
*PLT* expression in globular embryos (Kao et al., 2021). PLTs are clustered based on enrichment score.

**Table S1.** CRISPR mutants generated in this study. Mutation sites are indicated upstream of the PAM (N[GG] = 0) and downstream of the TSS (A[TG] = +1).

| Gene | Genotype | Indel | bp from PAM | bp from TSS | Length (aa) |
| --- | --- | --- | --- | --- | --- |
| *PLT2* | Col-0 | - | - | - | 568 |
|  | *plt2 bbm-cr* T<sub>1</sub> #1 | + T | -4 | +476 | 87 |
|  | *plt2 bbm-cr* T<sub>1</sub> #2 | + T | -4 | +476 | 87 |
|  | *plt2 bbm-cr* T<sub>1</sub> #3 | + A | -5 | +475 | 87 |
| *BBM* | Col-0 | - | - | - | 584 |
|  | *plt2 bbm-cr* T<sub>1</sub> #1 | + T | -4 | +21 | 15 |
|  | *plt2 bbm-cr* T<sub>1</sub> #2 | + A | -4 | +21 | 15 |
|  | *plt2 bbm-cr* T<sub>1</sub> #3 | + A | -4 | +21 | 15 |
|  | *plt3 bbm plt5 plt7-cr* | + A | -4 | +21 | 15 |

## Author contributions

V.W. designed the research. M.K., C.G., and V.W. performed the experiments. M.K., C.G., B.S. and V.W. wrote the manuscript. H.H. generated the promoter swaps. M.N. performed the transcriptome analysis. M.K. and C.G. contributed equally to the study. All authors have read and agreed to the published version of the manuscript.

## Acknowledgements

We thank Renze Heidstra for designing the sgRNAs used in this study.

## Funding

M.K. supported by GSGT.2019.019.

## Notes

### Competing Interest Statement

The authors have declared no competing interest.

